# Fitness-related traits are maximized in recently introduced, slow-growing populations of a global invasive clam

**DOI:** 10.1101/618082

**Authors:** Leandro A. Hünicken, Francisco Sylvester, Nicolás Bonel

**Author notes:** Author Contributions: LAH, FS, and NB conceived the study. LAH, FS, and NB conducted the literature search and extracted the data. NB designed the methodological approach. LAH and NB performed the data analysis. NB led the writing.

## Abstract

Many species are shifting their ranges being forced to rapidly respond to novel stressful environmental conditions. Colonizing individuals experience strong selective forces that favor the expression of life history traits notably affecting dispersal and reproductive rates in newly invaded habitats. Limited information is currently available on trait variation within the invasive range despite being critical for understanding ecological and evolutionary factors that drive the process of range expansion of invasive species. Here we evaluated life history shifts of the widely introduced Asian clam *Corbicula*, within its invaded range. Through an exhaustive literature search, we obtained data for 17 invasive *Corbicula* populations from different ecosystems worldwide. We tested the relationship between population and individual parameters relevant to the process of range expansion. Our main results are that recently introduced *Corbicula* populations were characterized by (i) low density and low rate of population increase, (ii) clams reproduced earlier in slow-growing populations, and (iii) density had no effect on population increase. All *Corbicula* populations analyzed in this study, which are fixed for one genotype (lineage Form A/R), experienced different selective environments in the introduced range. These findings support the perspective that adaptive phenotypic plasticity favored the expression of traits that maximize fitness in recently established populations, which faced stronger *r*-selective forces relative to long-established ones. We discuss the role of plasticity in facilitating rapid adaptation and increasing the likelihood of populations to overcome difficulties associated with low densities and low population increase in newly invaded areas.

## 1. INTRODUCTION

Climate change and global transport of people and goods is assisting the introduction of species worldwide, with associated economic and ecological impacts (Leppäkoski et al., 2002; Ruiz & Carlton, 2003; Walther et al., 2002). Hull fouling and ballast water associated to international shipping are among the strongest introduction vectors in place, and continue to introduce species into marine and freshwater ecosystems globally (Hewitt et al., 2009). Certain life history strategies facilitates the introduction and spread of invasive species in invaded areas (Zeeman et al., 2018). In order to prevent and mitigate the effects of species introductions, we need to better understand life history traits and strategies linked to species invasiveness. While an increasing number of studies on this topic is available, our understanding of processes that facilitate invasions is still incomplete (Rato et al., 2021). Comparisons among invasive populations of the same species could cast light into these processes, but such comparisons are rare.

Many species are forced to rapidly respond to novel stressful environmental conditions when shifting their distribution and colonizing new areas. Successful establishment and further spread of invasive species are influenced by both the abiotic and biotic nature of the colonized environment (Robinson et al., 2020; Soberon & Arroyo-Peña, 2017). Non-native species that have difficulties in adapting to their recipient environment, and fail to track abrupt environmental changes, become vulnerable to decline and extinction (Hill et al., 2011; Hoffmann & Sgró, 2011). Phenotypic plasticity is the ability of a single genotype to express different phenotypes when exposed to different environmental conditions (Travis, 1994; West-Eberhard, 2003). Plasticity provides the first step in adaptation to new environments on ecological time scales as long as costs to maintaining and/or producing a plastic response are not substantially large (Chevin & Lande, 2010; Ghalambor et al., 2007; Price et al., 2003). Adaptive plasticity responses favor the appearance of novel phenotypic variants, placing populations close enough to the new favored phenotypic optimum for directional selection to act (Ghalambor et al. 2007). These responses promote successful colonization and increase the likelihood for long-term persistence of invasive populations facing new environmental stress brought about by, for instance, range shifts (Hill et al., 2011; Pfennig et al., 2010; Pigliucci et al., 2006).

Colonization events through either continuous range expansion or jump dispersal are usually driven by a small number of individuals. While different range expansion mechanisms may be subjected to different selection forces and selection of characters, early colonizers benefit from increased space open for colonization and low conspecific density (Cole, 1954; Phillips et al., 2010; Sakai et al., 2001). Lag-times between initial colonization and the onset of rapid population growth are common in incipient, sparse populations (Sakai et al., 2001). The duration of the lag phase is mediated, among others, by a number of non-mutually exclusive ecological and evolutionary processes (e.g. low encounter rate for reproduction, density-dependent effects, adaptation, and selection of new genotypes; Bossdorf et al., 2005; Davis, 2005; Ricciardi, 2013). Following the lag phase, newly established populations encountering favorable conditions have the potential to grow exponentially (non-density regulated) while denser, long-established ones follow a density-regulated population growth (Brook & Bradshaw, 2006). According to theory, individuals at the expanding edge of the species distribution may experience strong *r*-selective forces (Burton et al., 2010; MacArthur & Wilson, 1967; Phillips, 2009). In favorable conditions, these forces can, for instance, favor the expression of traits that optimize population growth and survival (e. g. reduced age of maturity and reproduction, increased growth rate, increased energy allocated to reproduction; Stearns, 1976, 1977), thus affecting dispersal and reproductive rates (Phillips, 2009; Shine et al., 2011). An effective mechanism that increases the rate of population growth is early reproduction, which results from fast growing individuals (Cole, 1954; Lewontin, 1965; Roff, 1993). Individuals that grow faster will attain reproductive sizes earlier and bring down generation times, promoting a rapid build-up of population numbers (Cole, 1954; Lewontin, 1965; Phillips, 2009; Roff, 1993) and accelerating primary invasion and secondary spread of invasive species (Lockwood et al., 2007; Phillips et al., 2006; Shine et al., 2011; Skellam, 1951).

Several studies have compared life-history traits between native and invaded ranges (‘home-and-away’ comparisons). Such comparisons could lead to erroneous interpretations of the patterns observed because it would be difficult to disentangle what were the processes (e.g. propagule bias and/or evolution in populations at spatial (dis)equilibrium) causing the shifts in life-history traits (Phillips et al., 2010 and references therein). Interestingly, only a few studies have looked for such trait changes between introduced populations (Gaston, 2009), despite knowledge on life-history trait variation within the invasive range being critical for better understanding the process of species range expansion.

Bivalves of the genus *Corbicula* (Megerle Von Mühlfeld 1811) are native to Southeast Asia, the Middle East, Australia and Africa (Araujo et al., 1993) but have colonized a large part of the Americas and Europe (Crespo et al., 2015; Mouthon, 2001; Schmidlin & Baur, 2007). The invasion success of *Corbicula* clams has been mainly attributed to their rapid growth and maturation, high fecundity, and high dispersal rate (by gravid or potentially brooding adults; Prezant & Chalerwat, 1984). These characteristics make this bivalve one of the most successful invaders in aquatic ecosystems (McMahon, 2002; Sousa, Antunes, et al., 2008).

Invasive freshwater *Corbicula* populations are composed by a small number of lineages of hermaphroditic individuals reproducing through androgenesis (Komaru et al., 1998; Pigneur et al., 2012, 2014). The most widespread and abundant genotype in the invaded area is the lineage called Form A/R, widely known as *C. fluminea* (Pigneur et al., 2012, 2014). The current taxonomic status of the invasive *Corbicula* lineages in the Americas and Europe is still largely unresolved despite several morphological and genetic studies (reviewed in Pigneur et al. 2014). In the present paper, we deal with this form and refer to it as *Corbicula* to avoid entering unsettled taxonomic debates.

Here we investigated how different life-history traits of the Asian clam *Corbicula* responded to different selective pressures occurring at varying population contexts within the introduced range: (i) newly-vs. long-established, (ii) low vs. high density, and (iii) low vs. high rates of population increase; all of them being linked to the geographic range shifting of species’ distribution (i. e. front/core populations). To do so, we searched for peer-reviewed articles providing enough information to estimate time since population introduction as well as different population and individual processes and characteristics including population density, individual growth rate, minimum age at sexual maturity, and lifespan of this worldwide invader. We focused our predictions on life history strategies exhibited by individuals from recently established populations at the invasion front While site specific conditions (e.g., optimal vs. adverse) and spread history (e.g., continuous vs. jump dispersal) likely introduce differences among recently established populations, these are likely subjected to stronger *r*-selective pressures as compared to those from older, long-established ones. In the present work, we use a comprehensive dataset to capture interpopulation variability aiming to detect broad emerging patterns upon a global scale.

We tested the hypotheses that population density and population increase are directly related to time since introduction. *Corbicula* is widely known to exhibit rapid population growth, being able to reach extremely high densities (reviewed in McMahon, 2002). Different studies suggest that *Corbicula*’ s population dynamics are chiefly regulated by environmental conditions (e. g. Crespo et al., 2015, 2017; Gama et al., 2017), whereby population declines are typically driven by periodic mass mortality due to extreme hydrometeorological events (Ilarri et al., 2011; McDowell et al., 2017; McDowell & Sousa, 2019). We expected density to have no significant effect on population increase. Invasive *Corbicula* populations would follow a non-density regulated population increase, not leveling off at carrying capacity even at high densities as it happens with many other species (e.g. Brook and Bradshaw, 2006). Because high densities could negatively affect individual growth rates, we asked whether clams that grow comparatively faster and reproduce earlier are favored in slow-growing populations at the expanding front. A related prediction is that clams that grow rapidly would have a shorter lifespan with respect to individuals exhibiting slow growth.

## 2. METHODS

### 2.1 Literature search

We exhaustively searched the literature for peer-reviewed articles on *Corbicula* (*Corbicula fluminea sensu lato*) providing enough information to estimate the individual growth rate, population density, rate of exponential population increase, time since colonization, water temperature, and conductivity of global populations of this clam. To do this, we first identified relevant studies from ISI Web of Knowledge (Web of Science Core Collection, 1900 through 2017) by conducting a first search using the field tag “topic (TS)” in the “Advanced search” option and the search string: TS=“Corbicula fluminea” OR TS=“asia* clam” (search #1). We conducted a second search including topics TS=“population dynamic*” OR TS=“population size*” OR TS=“population densit*” OR TS=“population structure” (search #2). We ran a third search using TS=“ growth” OR TS=“individual growth” OR TS=“population growth” (search #3). Finally, we combined searches #2 and #3 into search #4 (#4= #3 OR #2), and restricted the latter by search #1 to obtain a final set of candidate studies (search #5= #4 AND #1).

This literature search was finished on 27 September 2018 and this protocol identified 215 candidate studies (database search: 215 candidates; gray literature and unpublished data: 0 candidates). To identify papers potentially missing in this list, we subsequently scanned the references in these papers and conducted a Google Scholar search using keywords mentioned above and added 8 studies to this candidate pool (Fig. S1). A total of 67 studies that belonged to research areas unrelated to the topic of the current study (*e.g*., applied chemistry, toxicology, paleontology) were excluded from this candidate pool by refining the set based in ‘Web of Science Categories’. The resulting studies (n = 156) were finally inspected individually to assess their eligibility for our analyses according to the following inclusion criteria: i) Studies that assessed invasive populations of *Corbicula* clams (i. e. studies exclusively providing estimates for native populations were excluded); ii) Studies that reported or provided enough data to extract the population density and age/size distribution for >6-months (see below); iii) Studies that assessed populations in reasonably “average” natural environments (populations from heavily impacted sites such as thermal plumes were excluded). We finally retained a sample of 16 studies, covering 17 invasive populations of *Corbicula* from different ecosystems worldwide (n = 140 out of 156 were excluded; see Fig. S1 for further details on screening and eligibility criteria). Whenever population data were only graphically presented, we contacted the authors directly. However, if data were no longer available or authors did not respond, we used the WebPlotDigitizer software (http://arohatgi.info/WebPlotDigitizer/) to extract data and accurately estimate population density and growth rates. This step-by-step criterion for the literature search and data collection attempted to reduce any potential selection bias in the long-term populations. This might occur because old high-density populations (with higher growth rates) were more frequently studied than young low-density populations (with low growth rates), which would be more likely to be extirpated if population characteristics were not changing with time.

### 2.2 Population density and exponential population increase

For each population selected from our literature search, we obtained the mean population density (ind/m^2^) for the time period considered in each study or extracted temporal values to estimate it, whenever this value was missing in the text. *Corbicula* clams generally exhibit a bivoltine juvenile release pattern (i.e. producing two generations a year McMahon, 2002; Sousa, Antunes, et al., 2008), leading to one or more high-density peaks related to recruitment events yearly. However, density in the field fluctuates almost constantly generating several peaks and valleys that do not necessarily indicate significant recruitment events. *Corbicula* populations exhibit rapid growth (Franco et al., 2012; McMahon, 2002; Sousa et al., 2006; Sousa, Antunes, et al., 2008). Hence, the rate of exponential population increase of *Corbicula* can be expressed by deriving the classical exponential growth equation:

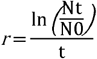

One of the goals of this study was to compare population growth rate during periods of substantial growth across populations to better understand *Corbicula* invasiveness. Hence, to estimate the rate of exponential population increase (*r*, hereafter population growth rate), we set a criterion to determine relevant recruitment peaks in each study by using the average density throughout the study period as a threshold. By doing so, we excluded time periods where population growth was extremely low, likely due to mass mortality, due to extreme hydro-meteorological events (e.g. a period of dramatic demographic decline). Likewise, this approach allows for exclusion of large time periods resulting from *Corbicula*’ s inability to reproduce year-round as they generally exhibit a uni-or bivoltine juvenile release. We therefore considered the lowest density value (*N*_0_), the highest peak above the average density (*N_t_*), and the time period elapsed between the valley and the peak (*t*). Intermediate density values comprised in each period (i. e. defined by *N_0_* and *N_t_*) were used to fit the growth equation and obtain *r*. Population rates of increase were estimated using GraphPad Prism version 7.00 for Windows, GraphPad Software, La Jolla California USA, www.graphpad.com. Periods defined by non-relevant peaks (i. e. below the average density threshold) were discarded as well as periods of progressive density decline between a relevant peak and the next *N*_0_ value (see Fig. S2 for further details on the step-by-step procedure used to estimate population increase). When populations exhibited more than one relevant density peak throughout the study period, the population increase rate for such populations was calculated as the average of single *r* estimations. Then, we standardized estimates on an annual basis to make them comparable. Note that the calculated *r* values provide a relative growth rate, that is, one that can be used to compare different populations to one another, rather than a population growth rate that can be used to forecast future population sizes.

### 2.3 Individual growth rates and parameters

For studies that only presented data on shell length-frequency distributions (shell length [mm] being the greatest distance between the anterior and posterior margins of the shell valve), we identified age cohorts using that information and applied the Bhattacharya’s method available in FISAT II software (Version 1.2.0, FAO-ICLARM Fish Assessment Tools; Gayanilo et al., 2002). To confirm each component of normal distributions from the modal progression analysis, we used the NORMSEP method also available in the FISAT II software (Pauly & Caddy, 1985). Once we had cohorts identified for all the studies, we calculated individual growth parameters using the same equations for all the studies considered. This procedure ensured that results were fully comparable across studies. For each cohort, in each of the populations considered in this study, we fitted the von Bertalanffy-growth function with seasonal oscillations (hereafter SVBGF) proposed by Pauly and Gaschutz (1979), Hoenig and Hanumara (1982), and Somers (1988):

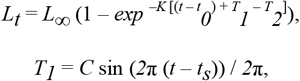

and

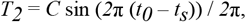

where *L_t_* is the predicted shell length at age *t, L*_∞_ is the asymptotic shell length, *K* is the growth constant of dimension time (year^-1^ in most seasonally oscillating growth curves) expressing the rate at which *L*_∞_ is approached, *t_0_* is the theoretical ‘age’ the clam has at shell length zero. *C* expresses the relative amplitude of the seasonal oscillation and varies between 0 and 1 (0 indicating lack of summer-winter differences in growth, it was constrained to be less than or equal to 1 during model fitting), and *t_s_* is the starting point of the oscillation. The parameters of the function were estimated by the modeling method available in JMP statistical software (v9.0 SAS Institute). When preliminary results of the SVBGF failed to converge to estimate the asymptotic shell length (*L*_∞_), we used the maximum shell length (*Lmax*) observed or reported at each study site to calculate the asymptotic shell length. To do this, we followed the equation suggested by Taylor (1958):

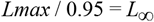

These values were fixed when we performed a second fit. Because the negative correlation between growth parameters (*K* and *L*_∞_) prevents making comparisons based on individual parameters (Ramón et al., 2007; Vakily, 1992), we used the growth-performance index (*GPI*), which reflects the growth rate of a given organism of unit length. In other words, *GPI* can be viewed as the (theoretical) value of *K* that would occur in organisms with an *L*_∞_ value of 1 unit of length (Munro & Pauly, 1983), and it was defined by Pauly and Munro (1984) as:

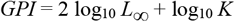

We calculated the *GPI* values for cohorts from each study site based on growth parameters (*L*_∞_ and *K*) obtained after fitting the SVBGF using the *Growth Performance Indices* application of the FISAT II software (Gayanilo et al., 2002).

### 2.4. Minimum age at sexual maturity

The minimum age at sexual maturity was estimated as the time (months) elapsed until clams reach the minimum size (shell length) at which invasive *Corbicula* clams become sexually mature, which, on average, is 10.0 mm ± 0.8 SE (n = 13 reported sizes). This value was obtained from shell sizes reported in the literature (Aldridge & McMahon, 1978; Cao et al., 2017; Eng, 1979; French & Schloesser, 1991; Gardner et al., 1976; Heinsohn, 1958; Ituarte, 1985; Kennedy & Van Huekelem, 1985; Kraemer, 1979; Kraemer & Lott, 1977; McMahon, 2002; Rajagopal et al., 2000). We acknowledge that in some conditions and populations, clams might mature at a different pace and thus introduce a degree of error in such cases. While this error is unavoidable until more detailed data is available, we are confident that the 10-mm estimate applies to many real-life situations and thus our results are robust.

### 2.5 Lifespan

As *t0* is the theoretical ‘age’ the clam has at shell length zero, *tmax* can be defined as the theoretical ‘age’ the clam reaches its maximum shell length (*Lmax*). Since *Lmax* is dependently related to a maximum point in time (years), it is assumed that *Lmax* implicates *tmax* (*Lmax* ⇒ *tmax*) (Bonel & Lorda, 2015). Thus, the theoretical maximum lifespan of a clam cohort can be approximately expressed as:

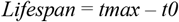

### 2.6 Time since colonization and key environmental variables

For the same 17 populations and sites studied, we calculated the time elapsed since population foundation (hereafter time since colonization) as the difference between the sampling dates in the first ecological survey and the first report of introduction. The accuracy of the introduction date is generally very difficult to know. However, many of the studies report range expansions of a conspicuous invader across well studied regions, where the appearance of the species might have been anticipated and readily detected by ongoing monitoring actions in the initial stages of invasion. Therefore, we consider the possible error introduced to be moderate to low and the value obtained as a good approximation. Moreover, we obtained water temperature and conductivity values for each site averaging monthly mean values for the same period used to estimate growth rates in each study. Other biotic and abiotic variables are known to interact with temperature to drive growth rates of invasive bivalves including food availability, dissolved oxygen, and heavy metal concentration (Bonel & Lorda, 2015; Vohmann et al., 2010). However, we only considered water temperature (°C) and conductivity (μS/cm) in the analyses because they have been reported to play a key role on the *Corbicula*’ s biology and ecology (Crespo et al., 2017; Franco et al., 2012; Magnussen, 2007). Indeed, temperature has emerged as the most important variable explaining the current distribution and, also, predicting that climate change will favor the expansion of *Corbicula*’s into colder areas at higher latitudes (Gama et al., 2017). Likewise, salinity has been considered a key factor influencing the success and velocity of the invasion of new estuarine ecosystems (e.g. Sousa et al., 2006). As *Corbicula* populations considered herein belonged to a broad range of freshwater and estuarine habitats, we therefore included conductivity to serve as a *proxy* for salinity levels. In order to keep values seasonally unbiased, only entire years were used in subsequent calculations. Whenever these parameters were not reported in the original paper, we obtained values for the same sites and years from the literature and reliable online databases (e. g. https://waterdata.usgs.gov/nwis/qw). When data was only graphically presented, we extracted it following the procedure mentioned above.

### 2.7 Statistical analyses

We ran five separate multivariate weighted linear regressions in order to test ecological processes and characteristics at three different (but non-mutually exclusive) levels: 1) population level, 2) population and individual level, and 3) individual level. In this sense, a response variable at the population level can be considered as an explanatory variable when analyzing a response at the individual level. In all cases, we computed the variance of the effect size from different populations, which were then used as weights in linear regressions. The inverse-variance weighting allows combining the results from independent measurements giving more importance to the least noisy measurements and *vice versa* (Borenstein et al., 2009).

We were interested in testing individual effects of predictors, which are related to lifehistory traits of *Corbicula* (hereafter reduced models). However, we added annual mean water temperature and conductivity as quantitative variables in each model (hereafter full models). Preliminary analyses showed that latitudinal effects are strongly correlated with water temperature (*r* = −0.84). Hence, it was excluded from the models to avoid multicollinearity. In some cases, we also included interaction terms between explanatory variables. We used the partial *F*-test to compare models—that is, full models with and without the interaction term, and full *versus* reduced models. As we had strong *a priori* hypotheses as to the direction of effects, we used one-tailed tests. Variables were Ln-transformed to fulfill the models’ assumption of normality and homoscedasticity, which were tested using the *gvlma* package (Peña & Slate, 2006). Prior to analyses, we standardized explanatory variables to zero mean and a unit of variance in order to get directly comparable effects. All analyses were carried out in R version 3.6.1 (R Development Core Team 2019). Values are given as means ± SE unless otherwise stated.

## 3. RESULTS

### 3.1 Literature search

Populations analyzed in this study belonged to Europe, North and South America (Fig. 1). These included eight populations in the United States, three in France, three in Argentina, and three in Portugal (see Table 1 for detailed location, bibliographic source, and code number used herein to refer to the populations, for each of the 17 *Corbicula* populations selected, and Table S1 for additional information).

**Figure 1.**
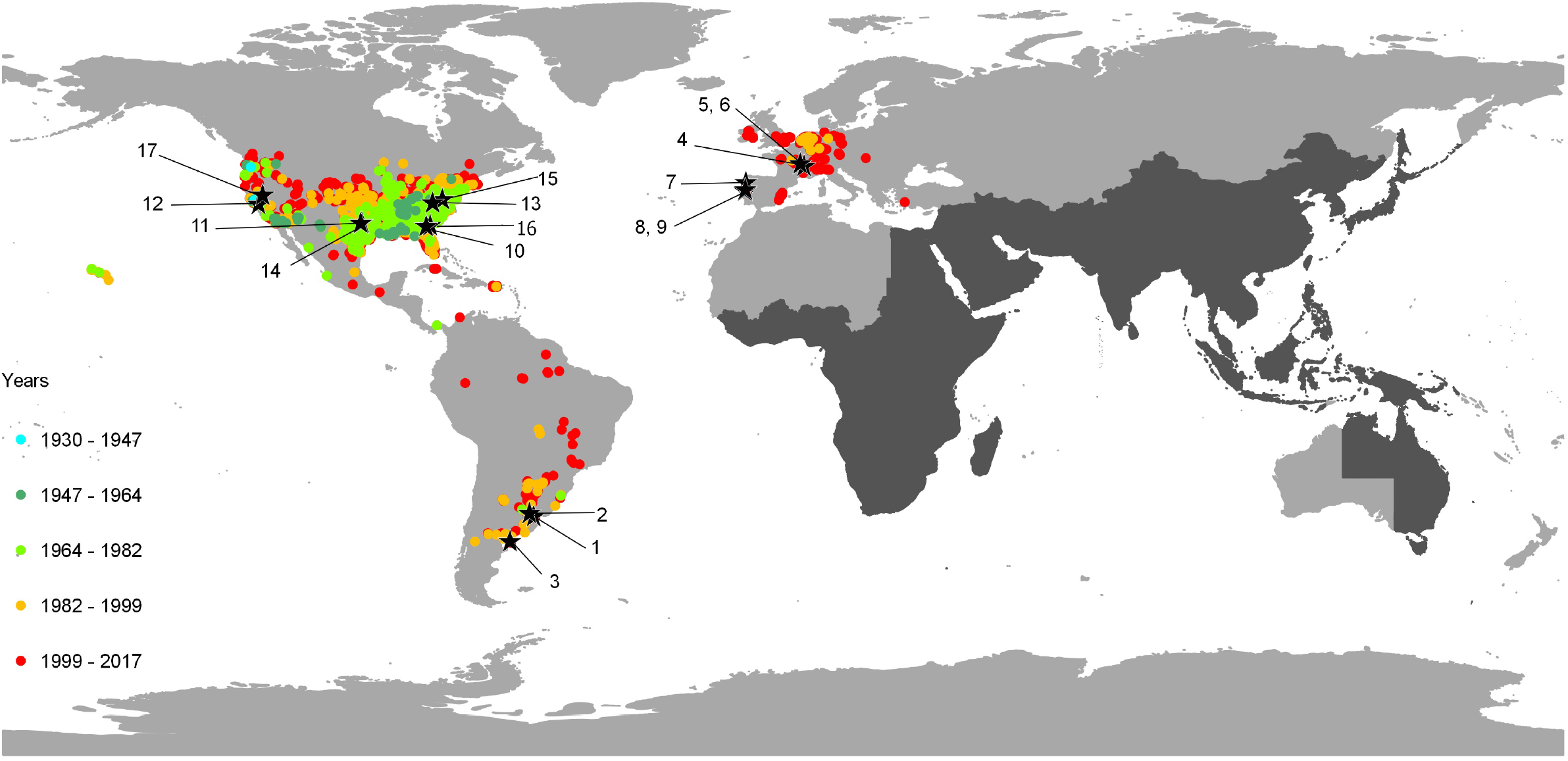
Global distribution of invasive *Corbicula* clams (Form A/R) indicating the native range of the genus (dark grey) and outside of the genus’ native range (color dots). Numbered stars are invasive populations reviewed in the present study and relates to ID# in Table 1. Years indicate estimated date of first introduction. References and details on the invasive populations are indicated in the mains text and Table 1.

**Table 1.** Summary of ecological and growth variables (mean ± SD) of worldwide invasive populations of the Asian clam *Corbicula* reviewed from the literature. *Density* is the estimated abundance of clams at the study site (ind./m^2^), *Pop.growth* is the rate of population growth, *Time* is the number of years elapsed since population colonization at the time of the original study (years), *Ind.growth* is the individual growth rate, *mASM* is the minimum age at sexual maturity (months), and *Lifespan* (years), *ID*# is a number assigned arbitrarily for identification purposes that relates to Figure 1. The country, specific study location, and reference to the source paper of the original study used to build the information are shown for each population. For extended information on each population and other environmental variables, see Table S2 of the Supporting Information.

| Country | Study location | Time | Population parameters |  | Individual parameters |  |  | ID# | Reference |
| --- | --- | --- | --- | --- | --- | --- | --- | --- | --- |
|  |  |  | Density | Pop.growth | Ind.growth | mASM | Lifespan |  |  |
| Argentina | Punta Atalaya, Río de la Plata | 3.5 | 339±405 | 2.20±0.01 | 2.87±0.07 | 4.30±0.13 | 3.83±0.57 | 1 | Ituarte (1985) |
| | Paraná de las Palmas River | 25.5 | 1,053±768 | 14.44±0.31 | 2.73±0.01 | 9.23±1.17 | 6.82±0.05 | 2 | Cataldo and Boltovskoy (1999) |
| | Río Negro estuary | 15.5 | 94±8.3 | 7.85±0.10 | 2.94±0.23 | 5.32±1.66 | 3.55±0.41 | 3 | Hünicken et al. (2019) |
| France | Saone River | 12.5 | 302±186 | 14.65±0.01 | 2.54±0.13 | 12.73±2.42 | 7.55±2.17 | 4 | Mouthon (2001) |
| | Loire Lateral Canal | 17.0 | 992±1209 | 41.17±0.12 | 2.80±0.24 | 8.93±5.95 | 5.17±3.00 | 5 | Mouthon and Parghentanian (2004) |
| | Canal of Roanne | 17.0 | 626±897 | 25.50±0.01 | 2.77±0.16 | 8.25±3.43 | 4.27±1.58 | 6 | Mouthon and Parghentanian (2004) |
| Portugal | River Minho estuary | 16.5 | 840±387 | 11.89±0.06 | 3.32±0.05 | 3.06±1.20 | 2.28±0.30 | 7 | Sousa et al. (2008b) |
| | Mondego estuary | 6.9 | 5,017±5,489 | 7.67±0.04 | – | – | – | 8 | Franco *et al.*, 2012 |
| | Casal de São Tomé | 10.3 | 3,255±1,115 | 4.06±0.04 | 3.14±0.12 | 3.83±1.69 | 2.75±0.51 | 9 | Rosa et al. (2014) |
| USA | Altamaha River | 4.5 | 712±1,613 | 0.37±0.00 | – | – | – | 10 | Gardner et al. (1976) |
| | Lake Arlington | 2.0 | 32±11 | 11.94±0.03 | 3.49±0.05 | 5.80±0.60 | 2.59±0.20 | 11 | Aldridge and McMahon (1978) |
| | Delta-Mendota canal | 28.0 | 9,470±7067 | 4.53±0.03 | 2.77±0.17 | 9.69±4.07 | 8.03±2.75 | 12 | Eng (1979) |
| | New River (Station 1) | 2 | 316±654 | 9.40±0.02 | – | – | – | 13 | Graney et al. (1980) |
| | Clear Fork, Trinity River | 7.0 | 4,474±3,899 | 19.34±0.04 | 3.00±0.16 | 6.76±3.62 | 7.07±2.83 | 14 | McMahon and Williams (1986) |
| | Mechums River | 10.5 | 669±636 | 188.52±0.62 | 2.56±0.15 | 9.20±4.41 | 3.31±1.21 | 15 | Hornbach (1992) |
| | Ogeechee River | 10.0 | 1,398±1,096 | 6.23±0.00 | 2.52±0.12 | 22.72±9.17 | 6.85±1.63 | 16 | Stites et al. (1995) |
| | Lake Tahoe | 8.0 | 1,809±723 | 16.64±0.04 | 2.80±0.12 | 6.90±1.90 | 2.70±1.32 | 17 | Denton et al. (2012) |

### 3.2 Density, growth, time since colonization, and key environmental variables

Overall, mean density of *Corbicula* from 17 locations worldwide spanned 32 to *ca.* 9,500 ind. m^-2^, population growth rate (*r*) varied from 0.37 to 189, individual growth rate (*GPI*) ranged 2.53-3.49, age at first reproduction 3.1-22.7 months, and lifespan 2.3-8.0 years (Table 1). On the other hand, time since colonization ranged 2 to 28 years in the populations studied (Table 1), yearly water temperature ranged from 13.2 to 20.7 °C, and conductivity from 68 to 8,457 μS/cm (see Table S1 of the Supporting Information for details). In all cases, we found no evidence that adding an interaction term with variables of interest, nor temperature and conductivity in each model, significantly increased its predictive power (for details see ‘*Model comparison*’ in Supporting Information). We therefore report estimates and statistics from the reduced models in Table 2.

**Table 2.** Summary of results from reduced models of weighted linear regression analyses of invasion and growth variables of global invasive populations of *Corbicula*. Parameter values are means ± SE. We report one-tailed *P* values indicating significant results in bold characters. Note natural logarithmic transformations of variables. For details on descriptive statistics of full models and results on partial *F*-tests see the Supporting Information. All variables were Ln-transformed and standardized to zero mean and a unit of variance (see *Statistical Analyses* for details).

| Response variable | Predictor | Estimate ± SE | Intercept ± SE | $R^2$ | *F*-statistics | num.df,<br>den.df | one-tailed<br>*P*-value |
| --- | --- | --- | --- | --- | --- | --- | --- |
| *Density* | *Time* | 0.454 ± 0.196 | -0.906 ± 0.241 | 0.263 | 5.34 | 1, 15 | **0.018** |
| *Population growth rate* | *Time* | 0.936 ± 0.323 | -0.518 ± 0.216 | 0.376 | 4.03 | 2, 14 | **0.006** |
| | *Density* | -0.522 ± 0.407 | | | | | 0.220† |
| *Individual growth rate* | *Density* | -0.422 ± 0.265 | 0.098 ± 0.264 | 0.283 | 2.17 | 2, 11 | 0.070 |
| | *Population growth rate* | -0.473 ± 0.344 | | | | | 0.099 |
| *minimum Age at sexual maturity* | *Individual growth rate* | -0.505 ± 0.179 | -0.192 ± 0.171 | 0.601 | 8.28 | 2, 11 | **0.008** |
| | *Population growth rate* | 0.460 ± 0.208 | | | | | **0.024** |
| *Lifespan* | *Individual growth rate* | -0.718 ± 0.184 | -0.056 ± 0.192 | 0.559 | 15.20 | 1, 12 | **0.001** |

† two-tailed *P*-value.

At the population level, we found that younger *Corbicula* populations (i.e. newly established) were characterized by low density and slow population growth rate with respect to older, long established ones (Fig. 2a, b). We found that density had no significant effect on population increase (Fig. 2c). When analyzing responses at the interaction between population and individual processes and characteristics, we found that clams tended to grow faster (marginally non-significant) in low-density and slow-growing populations (Fig. 2d, e). In contrast, we observed a strong positive response of the minimum age at sexual maturity as a function of population growth rate, clearly indicating that clams reproduce earlier in slow-growing populations (Fig. 2f). This is directly linked to processes occurring at the individual level, where clams with higher growth rates reach the minimum reproductive size (10 mm) earlier and have shorter life span than clams that grow more slowly (Fig. 2g, h).

**Figure 2.**
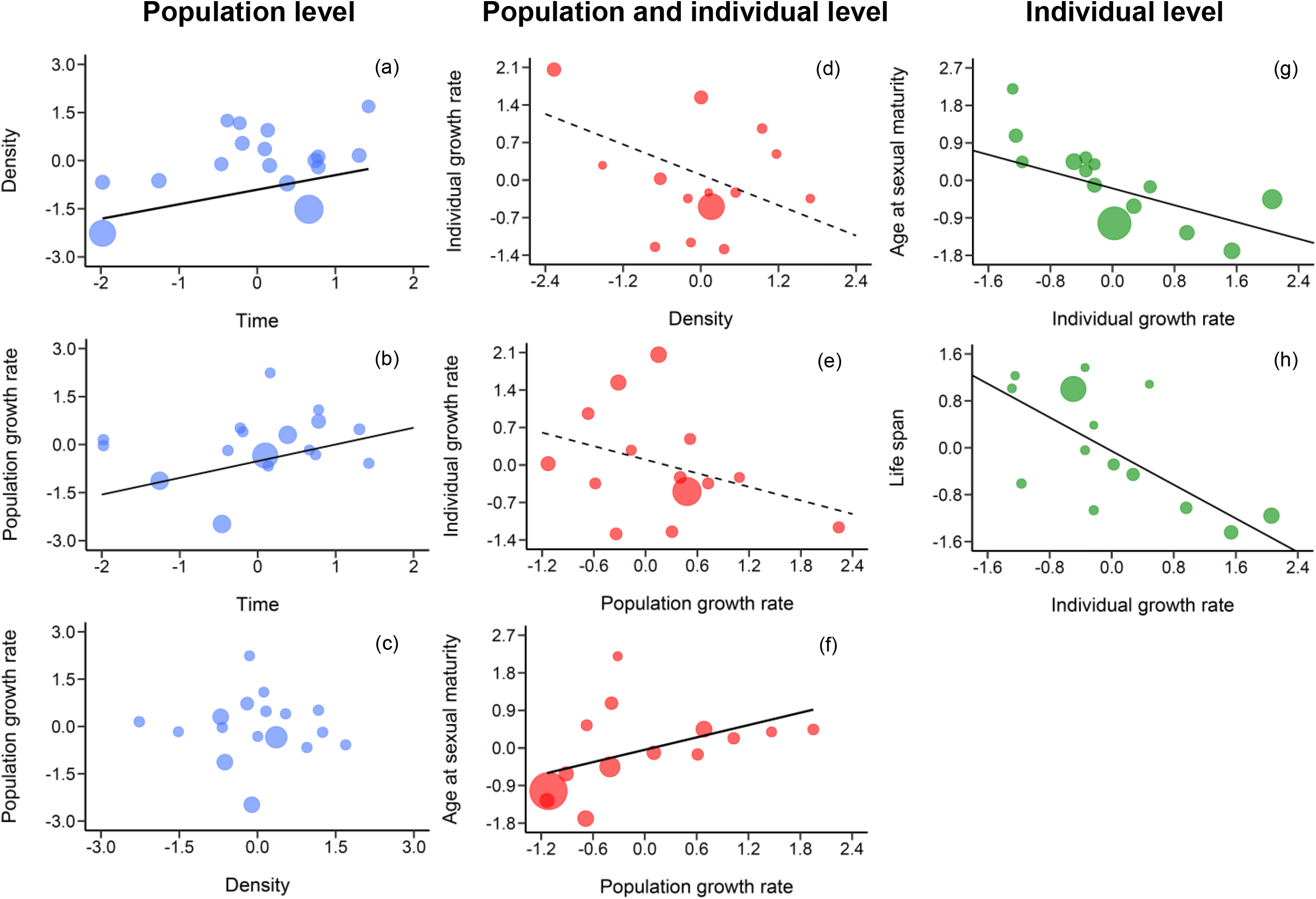
Relationships between processes and characteristics at different levels of organization of 17 global invasive populations of the Asian clam *Corbicula*. At the population level: a) the effect of time since colonization (*Time*, years) on population *Density* (ind./m^2^) and b, c) the simultaneous effect of *Density* and *Time* on *Population growth rate*. At the interaction between population and individual level: d, e) the simultaneous effect of density and population growth rate on individual growth rate, and f) the effect of *Population growth rate* on the minimum *Age at sexual maturity* (months). At the individual level: g) the effect of *Individual growth rate* on the minimum *Age at sexual maturity*, and h) on the *Life span* (years). When plotting the fit for one variable after testing for simultaneous effects, we removed the effect of the variable not considered (b, c, d, e). Dashed lines show fits when marginally non-significant 0.05> α <0.10. Solid lines show linear regression fits when statistically significant at α < 0.05. The size of circles correspond to the inverse of the variance of mean value of each observation of the response variable, used to weigh effect sizes in the regression analyses. All variables were Ln-transformed and standardized to zero mean and a unit of variance (see *Statistical Analyses* for details).

## 4. DISCUSSION

This is the first study that evaluates how population and individual parameters, involved in the process of range expansion, varied in response to different selective pressures occurring within the introduced range of the invasive Asian clam *Corbicula*. Our main results showed that recently introduced *Corbicula* populations were (i) characterized by low density and reduced population growth rate, (ii) clams reproduced earlier in slow-growing populations, and (iii) population growth rate was unfettered by density. These main findings are succinctly displayed in Figure 3.

**Figura 3.**
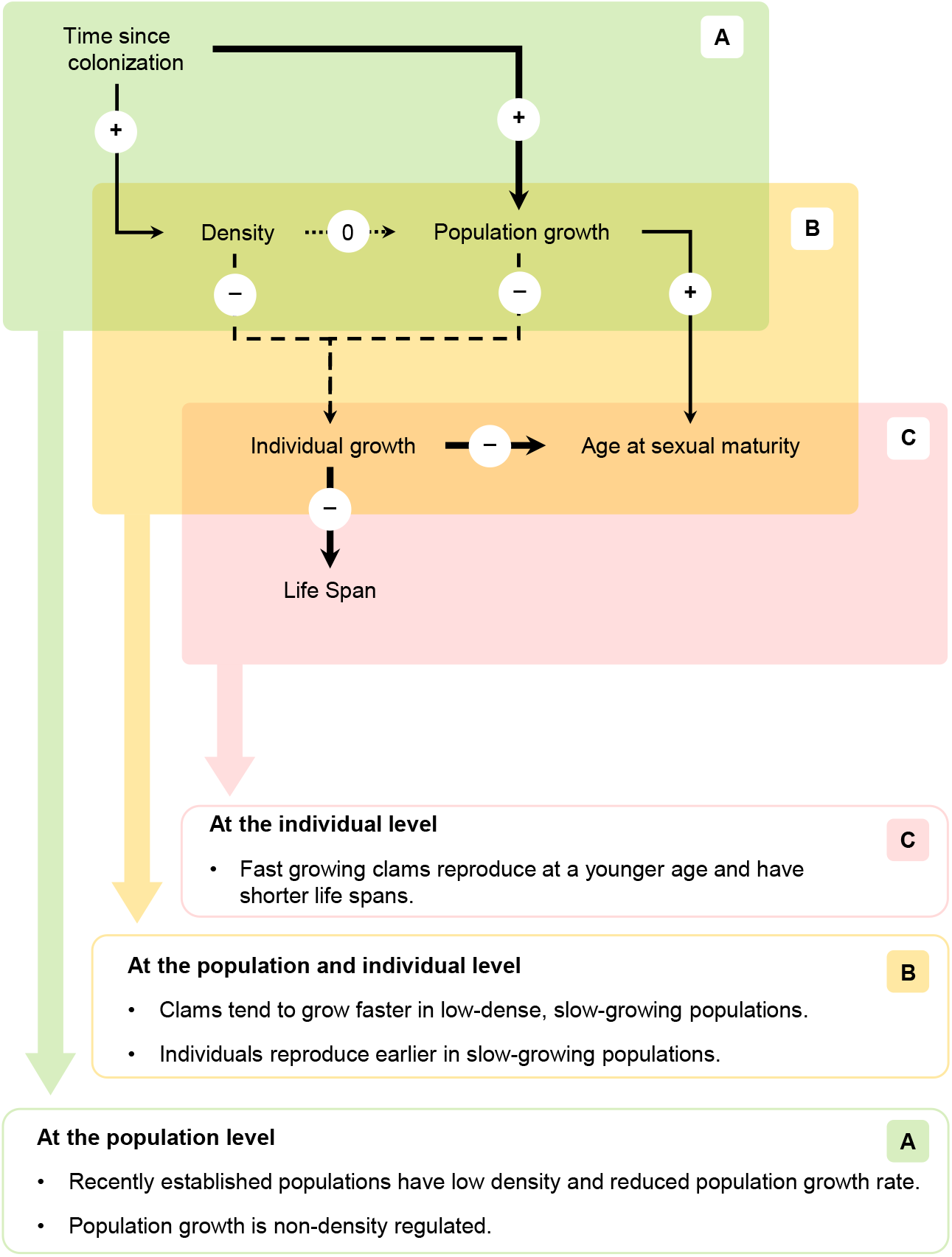
Summary of the relationships between processes and characteristics at different levels of organization of 17 global invasive populations of the Asian clam *Corbicula*, showing positive (+), negative (−), and no relationships (0) after fitting weighted linear regressions. Thick solid lines 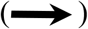 show linear regression fits when statistically significant at α < 0.01. Thin solid lines 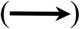 show linear regression fits when statistically significant at α < 0.05. Dashed lines 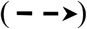 show fits when marginally non-significant 0.05> α <0.10. Dotted line 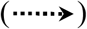 show no significant relationship α > 0.10. Relationships are shown at three different levels: A) at the population level, B) the interaction between processes and characteristics at the population and individual level, and C) at the individual level.

Density of *Corbicula* populations increased with time since colonization. This is consistent with the standpoint that recently established populations typically occur at conspecific density that is below carrying capacity (Cole, 1954; Sakai et al., 2001). Likewise, we found that population growth rate and time since colonization were directly related. This suggests that newly established populations experience stronger *r*-type selection that favors fitness-related traits and effectively increases the rate of population growth (Burton et al., 2010; MacArthur & Wilson, 1967; Phillips, 2009). Variation in population growth remained relatively low even at high-density values. Thus, population density did not predict changes in the rate of population increase. By definition, long established populations tend to be density regulated as they approach carrying capacity (logistic population growth; e. g. Phillips et al., 2010). Considering the so-called *r*/K selection continuum, this implies that newly established populations are predicted to shift from the *r*-towards the K-endpoint (MacArthur & Wilson, 1967). In other words, selection shifts a population from density-independent to density-dependent regulation. However, *Corbicula*’s population growth rate did not decrease with an increasing density even though some of the populations included in this study exhibit extremely high densities exceeding, in some cases, 100,000 individuals per square meter (e. g. Eng 1979).

Why was population growth unfettered by density regulation? Many studies have analyzed density dynamics of *Corbicula* and commonly assumed that, following exponential growth, the clam population achieves densities approaching the carrying capacity of the environment (e. g. McMahon, 2002; Sousa et al., 2006). However, to our knowledge, no studies have ever conducted field or laboratory experiments to test this hypothesis. There is a complete lack of knowledge on the true effect of density on population growth rate. Despite that a given *Corbicula* population might approach or even grow beyond the carrying capacity of the ecosystem, evidence from previous studies (and those cited herein) suggests that population size significantly varies with time. This means that the population is at non-equilibrium and that such fluctuations (or boom-bust dynamics; Strayer et al., 2017) are likely driven by density-independent factors. Indeed, *Corbicula*’ s dynamics are chiefly influenced by environmental variables such as water temperature, conductivity, pH, and/or dissolved oxygen, rather than by population’s density (e. g. Crespo et al., 2015; Gama et al., 2016). Periodic mass mortality due to extreme hydro-meteorological events as drought or extremely high or low temperatures could play an important role in infrequent, but catastrophic population declines (>99% in some cases; Ilarri et al., 2011; McDowell et al., 2017; Mouthon & Parghentanian, 2004; Sousa, Nogueira, et al., 2008). These events have been described (though not quantified) in several other systems as well (e. g. McDowell & Sousa, 2019; Vaughn et al., 2015). Thus, it does seem reasonable to infer that *Corbicula*’ s population growth rate is non-density-regulated and will not level off at carrying capacity. This supports the hypothesis that individuals from long established denser populations do not face a *K*-selective environment but a less strongly *r-*selective pressure.

Individuals tended to grow faster in low-density populations that exhibited low rates of population growth, shortening the time that clams need to reach maturity. As early-maturing clams reproduce earlier, this has the effect of promoting population increase for future population. The increase in individual growth was associated with a decrease in the clams’ life span likely due to trade-offs between life-history traits (Stearns, 1976, 1977). Such a trade-off is an effective mechanism to bring down generation times, which promotes a rapid population growth rate (Cole, 1954; Lewontin, 1965; Roff, 1993). In recently introduced populations, periods of negligible population increase are driven by non-mutually exclusive ecological and evolutionary adjustments such as the Allee effect, changing abiotic conditions, and adaptation and selection of new genotypes imposed by conditions in a new habitat (Bossdorf et al., 2005; Davis, 2005; Ricciardi, 2013; Sakai et al., 2001). This can lead to strong selection of life-history traits that optimize population increase and survival (Stearns, 1976, 1977). As predicted by life-history theory, individuals from recently established populations, occurring at the invasion front, face stronger *r-* selection relative to conspecifics from older populations (MacArthur & Wilson, 1967), favoring traits that accelerate the range expansion. For instance, some empirical observations have revealed that individuals from recently colonized areas grow faster, and thus, reproduce earlier, than individuals from older, long-established populations (e. g. butterflies: Hanski et al., 2006; fish: Masson et al., 2016; Australian cane toads: Phillips, 2009; tallow trees: Siemann & Rogers, 2001). Accordingly, our results support the perspective that recently introduced, slow-growing populations faced strong *r-*selection. In this way, incipient *Corbicula* populations increase their chances of overcoming difficulties associated with low densities and survive to the population establishment stage, contributing to the invasive success of this bivalve.

All invasive populations analyzed in this study are characterized by an extremely low genetic diversity and are fixed for one genotype (i.e. Form A/R; see Pigneur et al. 2012, 2014 for further details). Decrease in genetic diversity results in reduced evolutionary potential required to rapidly adapt to the novel habitat conditions of the invaded range, among other issues related to critical invasion phases (e.g. initial colonization, leading edges of range expansion; Schrieber & Lachmuth, 2017). The question that inevitably arises is how genetically depleted *Corbicula* populations succeed in their invasion process. This poses a dilemma known as the genetic paradox of invasion (Allendorf & Lundquist, 2003). A growing number of studies propose different mechanisms that allow the invasive population to increase population fitness and compensate the loss in genetic diversity during introduction events and in the invasion front (see review by Estoup et al., 2016 for further details). Adaptive phenotypic plasticity is one possible mechanism that could compensate genetic diversity loss and explain high invasiveness of *Corbicula* populations. Specifically, plasticity favors the appearance of novel phenotypic variants in an introduced area, placing populations close enough to the new phenotypic optimum despite potentially lower genetic variation (Estoup et al., 2016; Ghalambor et al., 2007). The shift in fitness related traits provides a reasonable ground for advocating that adaptive plasticity facilitated the expression of phenotypic variants among *Corbicula* populations experiencing different environmental circumstances. Further studies would require an appropriate quantitative approach to measure other fitness components to determine which of the strategies encompassed by adaptive plasticity is followed by invasive *Corbicula* clams (see examples: Davidson et al., 2011; Richards et al., 2006).

In conclusion, our results showed that environmental differences, linked to varying population contexts, generated strong selective pressures that induced important shifts in the expression of life-history traits of the Asian clam. Individuals from recently established, slow-growing populations grew faster and thus reproduced earlier than those from long-established ones. These findings support the perspective that adaptive plasticity favored the expression of traits that maximize fitness in response to stronger *r*-selective forces, thus decreasing the risk of stochastic extinction associated with low densities at the invasion front. Population growth rate was unfettered by population density and seems unlikely to level off at carrying capacity. This suggests that *Corbicula* populations are at non-equilibrium and that its dynamics is chiefly influenced by environmental variables rather than density-dependent factors. Other relevant variables could not be considered in our models. However, this study represent a first step in the attempt to understanding how life-history traits of the Asian clam respond to different selective regimes in the introduced range. Such information is critical to better comprehend the processes and mechanisms promoting rapid adaptation and thus successful colonization of invasive populations facing new environmental stress brought about by range shifts.

## Notes

### Competing Interest Statement

The authors have declared no competing interest.

### Summary of Updates

In the present version, we have revised the entire manuscript and associated files, thoroughly modified them, conducting new analyses, modifying/including new figures, and rewriting full sections as necessary

