## Supporting Information for "Fitness-related traits are maximized in recently introduced, slow-growing populations of a global invasive clam"

**Table of Contents**

***Supplementary Tables***

*Table S1. Summary of model comparison*	2

*Table S2. Summary of means of variables estimated for* Corbicula *populations*	3

***Supplementary Figure***

*Fig. S1. PRISMA diagram for literature search*	4

*Fig. S2. Steps for estimating population increase*	5

**Supplementary Tables**

**Table S1.** Summary of the *F*-tests after model simplification between full and reduced weighed linear models at different levels of organization. *Density* is the estimated abundance of clams at the study site and for each studied period (ind./m2), *Pop.growth* is the rate of exponential population derived from the classical form of the exponential growth equation, *Time* is the number of years elapsed since population colonization at the time of the original study (years), *Ind.growth* is the individual growth rate (GPI, a proxy for individual growth rate), *mASM* is the minimum age at sexual maturity (months), *Lifespan* (years), *Temperature* is the mean annual water temperature (ºC) of the reported location (numbers in parentheses indicate minima and maxima estimates), and *Conductivity* is the mean annual conductivity (µS/cm) of the reported location. All variables were Ln-transformed and standardized to zero mean and a unit of variance (see *Statistical Analyses* for details).

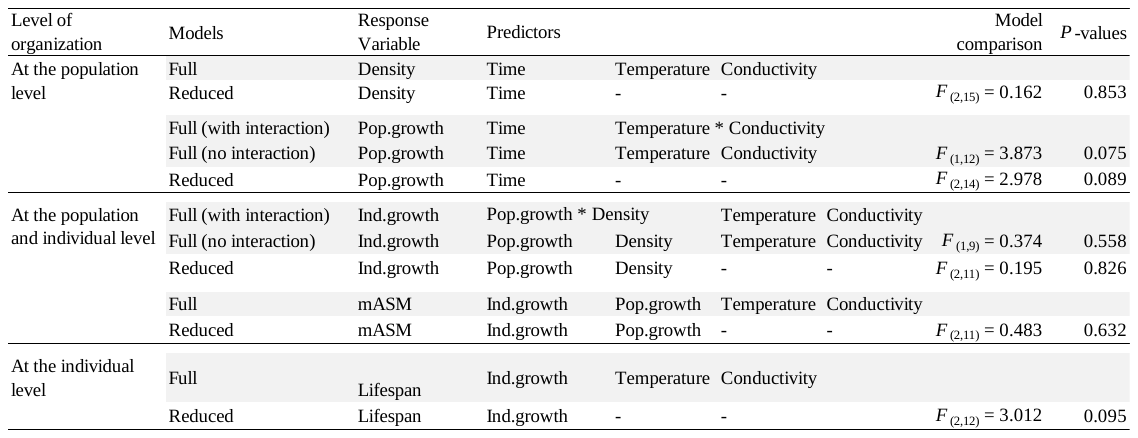

**Table S2.** Summary of ecological and growth variables (mean ± SD) of worldwide invasive populations of the Asiatic clam Corbicula reviewed from the literature. *ID*# is a number assigned arbitrarily for identification purposes, *L*∞ is asymptotic length (mm), *K* is the growth constant (year^-1^), *C* is the relative amplitude of the seasonal oscillation, *Temp* is the mean annual water temperature (ºC) of the reported location (numbers in parentheses indicate minima and maxima estimates), *Cond* is the mean annual conductivity (µS/cm) of the reported location, *Lat* and *Long* are the latitude and longitude of the study sites, *Year reported* indicates the first report of the clam and *Year study* indicates the year when the first ecological survey was done; both dates were used as a proxy to estimate the time elapsed since the population colonization. NA, data not available.

| Country | Study location | *ID#* | *L∞* | *K* | *C* | *Temp* | *Cond* | *Lat* | *Long* | *Year reported* | *Year study* | References |
| --- | --- | --- | --- | --- | --- | --- | --- | --- | --- | --- | --- | --- |
| Argentina | Punta Atalaya, Río de la Plata | 1 | 31.3 | 0.77±0.12 | 0.88±0.17 | 19.4 (11.0-27.0) | 647 | -35.01 | -57.54 | 1982 | 1985 | Ituarte (1985) |
|  | Paraná de las Palmas River | 2 | 34.7 | 0.44±0.01 | 0.75±0.35 | 18.2 (11.9-28.1) | 121 | -34.30 | -58.52 | 1970 | 1996 | Cataldo & Boltovskoy (1999) |
|  | Río Negro estuary | 3 | 34.6±1.7 | 0.83±0.34 | 0.90±0.15 | 15.0 (6.1-24.6) | 304 | -40.80 | -63.03 | 1997 | 2013 | Hünicken et al. (2019) |
| France | Saone River | 4 | 28.9 | 0.42±0.11 | 1.00±0.00 | 13.2 (1.0-24.9) | 506 | 45.80 | 4.84 | 1985 | 1998 | Mouthon (2001) |
|  | Loire Lateral Canal | 5 | 31.0 | 0.73±0.33 | 1.00±0.00 | 14.9 (2.8-24.9) | 232 | 46.49 | 3.87 | 1985 | 2002 | Mouthon & Parghentanian (2004) |
|  | Canal of Roanne | 6 | 28.5 | 0.76±0.27 | 1.00±0.00 | 14.3 (2.7-24.0) | 213 | 46.33 | 4.00 | 1985 | 2002 | Mouthon & Parghentanian (2004) |
| Portugal | River Minho estuary | 7 | 38.8±3.62 | 1.41±0.16 | 0.75±0.12 | 22.2 (1.0-24.9) | 98.3 | 42.05 | -8.60 | 1989 | 2006 | Sousa et al. (2008b) |
|  | Mondego estuary | 8 | NA | NA | NA | 16.4 (10.5-24.2) | 8457 | 40.13 | -8.83 | 2001 | 2008 | Franco et al. (2012) |
|  | Casal de São Tomé | 9 | 33.7 | 1.26±0.37 | 0.93±0.16 | 14.9 (11.2-19.8) | 382 | 40.42 | -8.74 | 2002 | 2012 | Rosa et al. (2014) |
| USA | Altamaha River | 10 | NA | NA | NA | 20.0 (6.5-30.0) | 76.9 | 31.94 | -82.34 | 1969 | 1973 | Gardner et al. (1976) |
|  | Lake Arlington | 11 | 49.1 | 1.28±0.14 | 1.00±0.00 | 20.7 (10.0-33.0) | 270 | 32.72 | -97.19 | 1973 | 1975 | Aldridge & McMahon (1978) |
|  | Delta-Mendota canal | 12 | 40.4 | 0.39±0.16 | 0.97±0.06 | 18.4 (8.4-25.3) | 563 | 37.07 | -121.00 | 1945 | 1973 | Eng (1979) |
|  | New River (Station 1) | 13 | NA | NA | NA | 13.9 (1.5-29.1) | 116.1 | 37.38 | -80.85 | 1975 | 1977 | Graney et al. (1980) |
|  | Clear Fork, Trinity River | 14 | 47.2 | 0.47±0.15 | 1.00±0.00 | 19.9 (8.9-29.4) | 314 | 32.67 | -97.44 | 1974 | 1981 | McMahon & Williams (1986) |
|  | Mechums River | 15 | 20.0 | 0.94±0.32 | 0.69±0.33 | 26.7 (10.5-40.8) | 68 | 38.15 | -78.60 | 1972 | 1983 | Hornbach (1992) |
|  | Ogeechee River | 16 | 27.4 | 0.46±0.12 | 0.79±0.15 | 15.9 (0.0-26.4) | 114 | 32.14 | -81.41 | 1974 | 1984 | Stites et al. (1995) |
|  | Lake Tahoe | 17 | 23.2 | 1.31±0.46 | 0.98±0.03 | 14.2 (5.4-20.0) | 78 | 39.13 | -120.05 | 2002 | 2010 | Denton et al. (2012) |

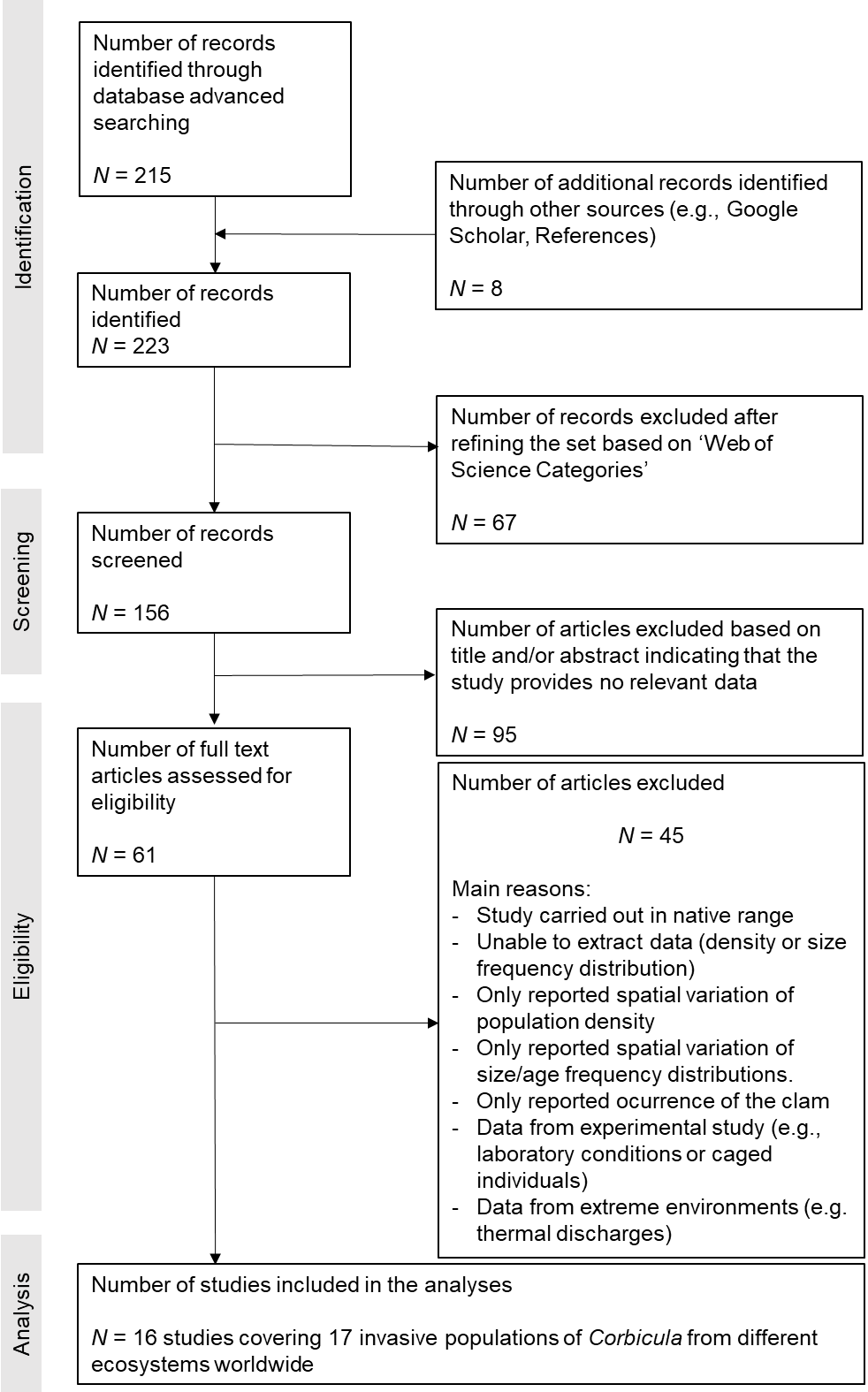
**Supplementary Figures**

**Figure S1.** PRISMA diagram of data flow and paper selection showing exclusion criteria across different phases of the literature search for *Corbicula* (*Corbicula fluminea sensu lato*) peer reviewed papers to be analyzed in the present study. See main text for search details. (Modified from Janicke et al. 2016.)

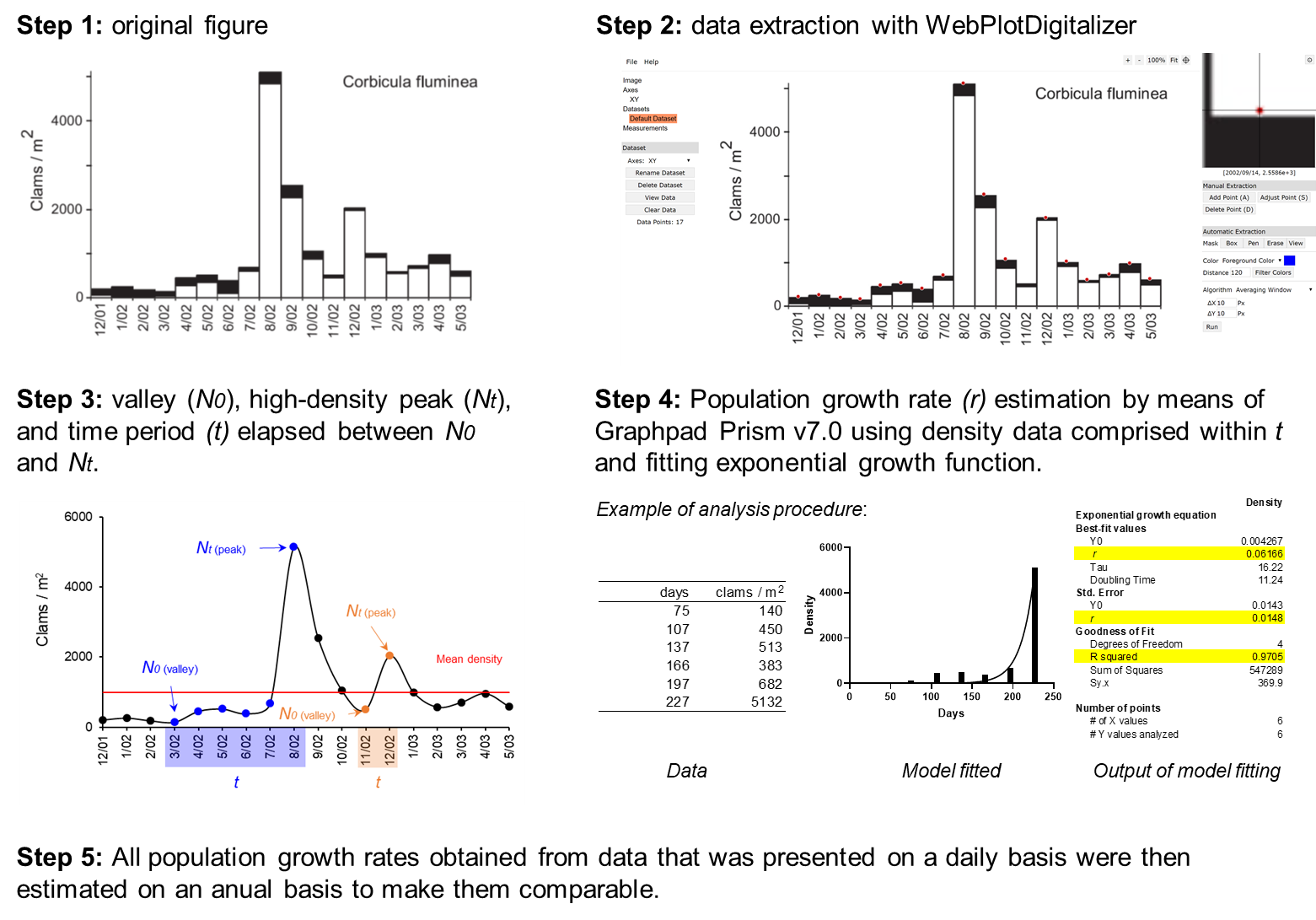

**Figure S2.** Step-by-step procedure to estimate population increase from data collected from literature where mean density was used as threshold to define peaks (*Nt*), valleys (*N_0_*), and the time period (*t*) between them.
